# Genetically edited human placental organoids cast new light on the role of ACE2 in placental development

**DOI:** 10.1101/2024.05.09.592870

**Authors:** Anya L. Arthurs, Bianca Dietrich, Martin Knöfler, Caleb J. Lushington, Paul Q. Thomas, Fatwa Adikusuma, Tanja Jankovic-Karasoulos, Melanie D. Smith, Kirsty G. Pringle, Claire T. Roberts

## Abstract

ACE2 expression is altered in pregnancy disorders and *ACE2* gene variants are associated with several major pregnancy complications including small-for-gestational-age, fetal growth restriction and preeclampsia. This study utilised gene-editing to generate both *ACE2* knockout and *ACE2* rs2074192 placental organoids, facilitating mechanistic studies into the role of *ACE2* in placental development, and the effect of fetal carriage of *ACE2* rs2074192 CC, CT and TT genotypes. Parameters of cell and organoid growth were measured, together with qPCR, Western Blotting, and ELISA assessments, in all groups from both organoid models. Here, we report that *ACE2* knockout results in delayed placental cell growth and increased cell death. *ACE2* knockout organoids had lower ACE protein expression, reduced organoid diameters and asymmetrical growth. Placental organoids with the ACE2 rs2074192 TT genotype had significantly higher expression of *ACE2* mRNA and ACE2 protein with elevated ACE2:ACE expression ratio and no change in ACE protein expression. Despite increased expression of ACE2 protein, ACE2 enzyme activity was significantly decreased in ACE2 rs2074192 TT placental organoids. TT organoids also had reduced diameters and asymmetrical growth. Our research provides new molecular understanding of the role of ACE2 in placental development, with potential implications for pregnancy in carriage of the *ACE2* rs2074192 gene variant.

## Introduction

Understanding the molecular mechanisms underlying pregnancy and fetal development is crucial for improving maternal and neonatal health outcomes. Despite significant advances in obstetric care, complications such as small-for-gestational-age, fetal growth restriction and preeclampsia continue to pose substantial risks. Recent studies have highlighted the pivotal role of placental function in these conditions, with literature suggesting that variations in placental angiotensin converting enzyme 2 (*ACE2*) gene expression may be a key factor. However, there remains a gap in our comprehensive understanding of how *ACE2* genetic variations and molecular pathways contribute to these pregnancy disorders.

Interest in previously little-known ACE2 has escalated with the COVID-19 pandemic and the knowledge that ACE2 is the SARS-CoV-2 receptor. However, the role of ACE2 in the placenta remains understudied, despite its evident importance in pregnancy. Pregnant women have higher circulating levels of ACE2 compared to non-pregnant women^1^. ACE2 abundance is altered in pregnancy disorders, including preeclampsia, small for gestational age (SGA) and fetal growth restriction^1,2,3,4,5,6,7^, and *ACE2* gene variants are associated with several major pregnancy complications^1, 5, 8^. Genetic factors account for two thirds of the phenotypic variation in circulating ACE2^9^; specifically, *ACE2* single nucleotide polymorphisms (SNPs) and haplotypes are associated with altered abundance of Angiotensin II (Ang II) and Angiotensin 1-7 (Ang 1-7), which are the peptides produced by angiotensin converting enzyme (ACE) and ACE2 catalytic activity, respectively^10^ (**Figure 1**).

**Figure 1.**
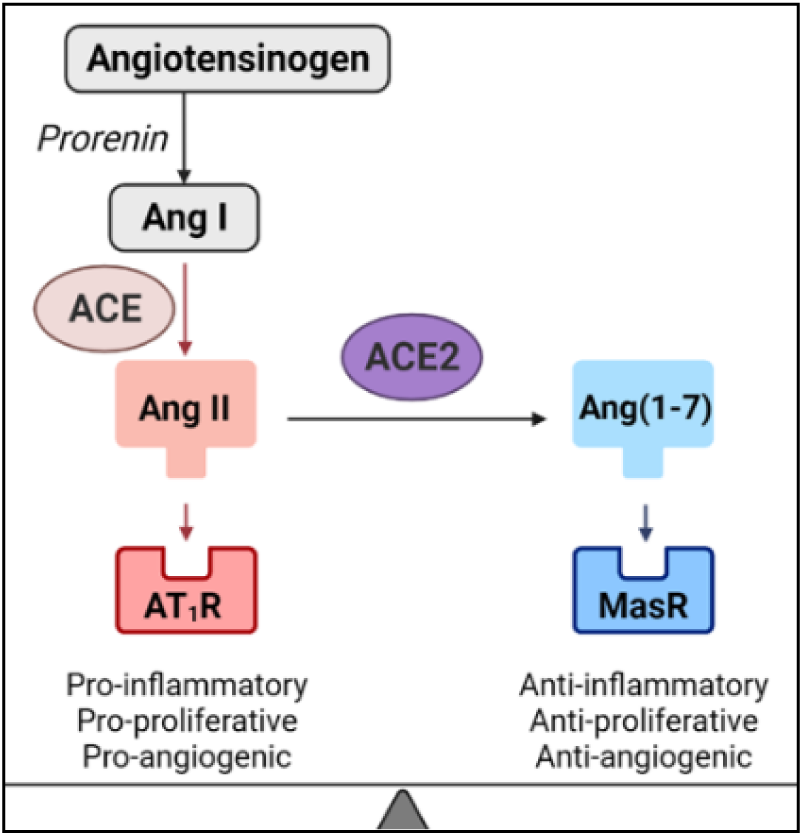
The ACE- and ACE2-mediated opposing arms of the renin-angiotensin system.

The circulating renin-angiotensin system is a critical regulator of blood volume and systemic vascular resistance. However, in addition to the circulatory system, local tissue-specific renin-angiotensin systems exist, including within the placenta. The pro-inflammatory, pro-proliferative arm of the placental renin-angiotensin system (shown in red in **Figure 1**), mediated by ACE, promotes cell and tissue growth. This is opposed by an anti-inflammatory, anti-proliferative arm (shown in blue in **Figure 1**), mediated by ACE2^3^. ACE and ACE2 maintain a finely tuned balance within the renin-angiotensin system to optimise placental growth and function throughout healthy pregnancy. However, the expression and activity of ACE and ACE2 enzymes, both systemically^1, 2^ and in the placenta^3,4,5,6,7^, is disturbed in many pregnancy complications associated with renin-angiotensin system dysfunction.

Adequate ACE2 expression and activity, in both the mother and placenta, is necessary for healthy pregnancy. In *ACE2^-/-^*(*ACE2* knockout) mouse models, *ACE2* knockout dams have higher systolic blood pressure^2, 4^, decreased plasma Ang 1-7^2^, and placentae have increased levels of^2^, and sensitivity to^4^, Ang II. This results in alterations to the balance of ACE and ACE2 activity in the renin-angiotensin system. Furthermore, pups from *ACE2* knockout dams have reduced weight^2, 4^ and length^2^, with lower pup to placenta weight ratios, indicating compromised placental function. Clearly ACE2 is essential for healthy pregnancy but dysregulation of ACE2 levels and activity, as well as ACE:ACE2 balance, is pathogenic^4, 11^. However, whilst murine research is informative, an *ACE2* knockout model of human placenta has never been established.

As previously mentioned, *ACE2* gene variants (including single nucleotide polymorphisms [SNPs]) alter expression of renin-angiotensin system components. A SNP is a single base substitution at a specific chromosome location. Many SNPs in the *ACE2* gene have been associated with disease states^5, 8, 12,13,14^. In particular, *ACE2* rs2074192 is associated with increased incidence of diseases characterised by renin-angiotensin system dysregulation^12,13,14,15^, including an increased susceptibility to COVID-19 and increased disease severity^13, 14^. Importantly for pregnancy, SGA, a condition where neonatal birthweight is lower than expected for their gestational age, has been associated with *ACE2* rs2074192 (C to T polymorphism) which, when carried by the fetus, increases the risk of SGA by 23-fold^5^. The placenta, a product of conception, is genetically identical to the fetus. As such, a fetal SNP will also be present in the placenta. Despite strong associations between *ACE2* rs2074192 frequency and disease state, and the estimated frequency of at least one copy of this SNP in one third of the global population^16^, research is mostly limited to association studies but not its mechanisms of action.

In this study, we utilised gene-editing to generate *ACE2* knockout placental organoids and *ACE2* rs2074192 placental organoids, creating the first induced SNP gene-edited organoids in human placenta. We investigated the role of *ACE2* in placental development by profiling expression of *ACE2* mRNA and protein, ACE protein, ACE2 enzyme activity and parameters of cell and organoid growth in both placental organoid models. Elucidating the role of *ACE2* in the placenta will allow inferences to be made regarding its overall contribution to pregnancy health, as well as its possible role in other physiological and pathological states that involve the renin-angiotensin system.

## Methods

Trophoblast stem cells (TSC) were isolated from n=3 patients and grown to confluence. At this point, TSCs from each patient were separated into two groups:

*1.* *ACE2^-/-^* (*ACE2* knockout – hereafter referred to as *ACE2^+/+^, ACE2^+/-^* or *ACE2* KO)
*2.* *ACE2* SNP (rs2074192 – hereafter referred to by genotype, i.e. CC, CT or TT)

The generation of gene-edited organoids from each patient was replicated three times, thus a final n=9 independently generated organoids for each genotype group (see **Figure 2**).

**Figure 2.**
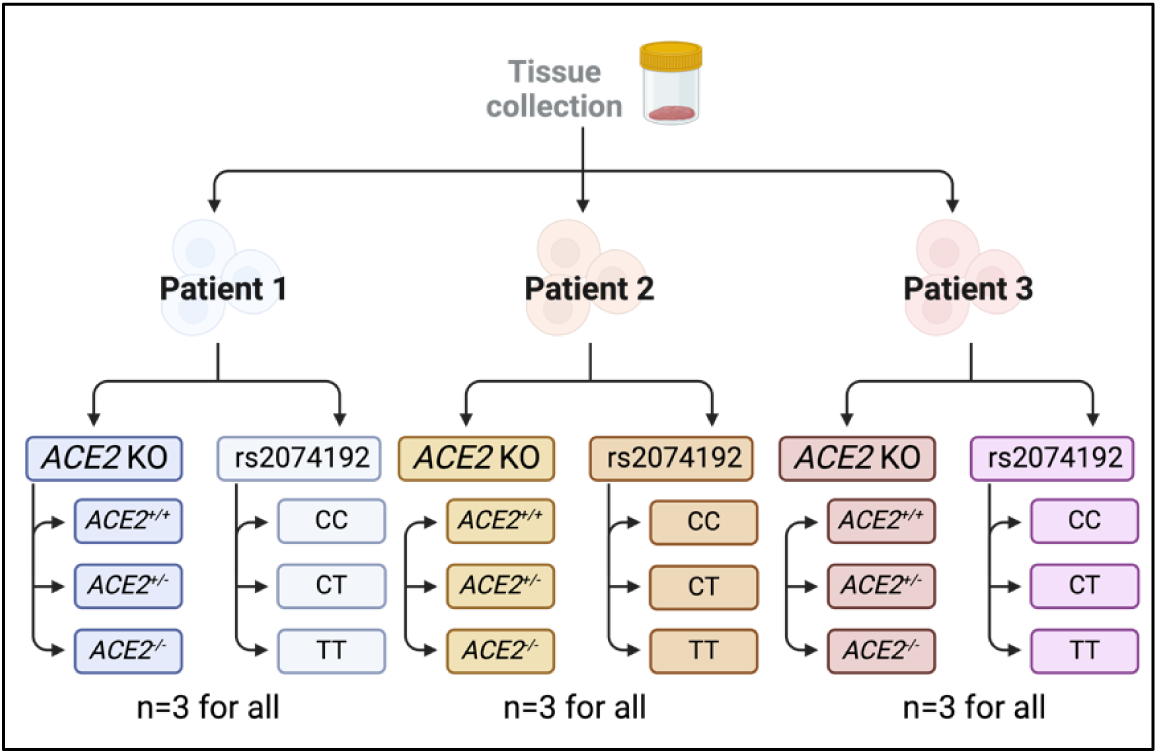
Method describing gene-editing to generate different TSC groups.

As *ACE2* is on the X chromosome, only placentae from female fetuses were included to allow analysis of a heterozygous group (*ACE2^+/-^*).

### Tissue collection

First trimester (6-7 weeks’ gestation) human placentae were obtained with informed consent from women undergoing elective termination of pregnancy at the Pregnancy Advisory Centre in the Queen Elizabeth Hospital, in Woodville, South Australia. Ethics approval was obtained from the Central Adelaide Local Health Network Human Research Ethics Committee, HREC/16/TQEH/33, Q20160305. Placentae were collected within minutes of termination upon which villous tissue was washed with Hanks’ Balanced Salt Solution (HBSS; Gibco, Sigma-Aldrich) and transported to the laboratory on ice.

### Genotyping for fetal sex

Placentae from n=5 patients were genotyped for fetal sex high resolution melt curve analysis of the gene that encodes amelogenin. Amelogenin is found on both the X (AMELX) and Y (AMELY) chromosomes, the X allele features a 3 bp deletion in exon 3 allowing the identification of samples with only X chromosomes (female) or X and Y chromosomes (male). Forward primer 5ʹ-CCCTGGGCTCTGTAAAGAATAGTG-3ʹ, reverse primer 5ʹ-ATCAGAGCTTAAACTGGGAAGCTG-3. qPCR was performed using SSOFast EvaGreen Supermix (Bio-Rad, CA), primers at 250 nM final concentration, 5 ng of DNA per reaction, on a Bio-Rad CFX384 Real-Time PCR System. Cycling conditions: initial denaturation 98°C 30 s, 40 cycles of 98°C for 5 s and 60°C for 5 s. High resolution melt curve analysis was performed from 65°C to 85°C with a 0.2°C increment every 10 s. Melt curve between 65°C and 70°C was analysed using Bio-Rad Precision melt software (Bio-Rad, CA) to identify sex genotypes. N=3 of these placentae, determined to be from female fetuses, were selected for use in this study.

### Isolation and cultivation of TSCs

TSCs were isolated from first trimester placentae according to Dietrich, *et al*.^17^ Briefly, first trimester villous tissue was washed in HBSS (4°C) prior to manually isolating villus structures for further processing. Villi in HBSS were centrifuged (1000 rpm, 1 min) prior to three consecutive enzymatic digest steps and density gradient centrifugation as per Haider, *et al.*^18^. After careful collection of cells between the 35% and 50% Percoll layers, cells were thoroughly washed with HBSS before seeding in fibronectin-coated culture plates using a culture medium to promote stemness. As per Dietrich, *et al.*^17^, TSC medium contained DMEM/F12 (Gibco) supplemented with 1×B-27 (Gibco), 1×Insulin-Transferrin-Selenium-Ethanolamine (ITS-X; Gibco), 1 µM A83-01 (Tocris), 50 ng/ml recombinant human epidermal growth factor (rhEGF; R&D Systems), 2 µM CHIR99021 (Tocris) and 5 µM Y27632 (Santa Cruz).

### Generation of plasmids

To generate modified ACE2 organoid models, two distinct gene editing strategies were employed. Firstly, a modified pDG459 plasmid (Addgene #100901) was designed to induce a complete *ACE2* knockout (KO). This entailed the precise targeting of regions approximately 100 base pairs upstream and downstream of the first and last *ACE2* exons, resulting in the excision of promoter, regulatory, and coding sequences. Secondly, a modified PEA1-Puro plasmid (Addgene #171991) was designed to introduce an intronic *ACE2* single nucleotide polymorphism (SNP), specifically rs2074192. The SNP was targeted using the PE3 strategy, which incorporated a downstream nicking single-guide RNA (sgRNA) positioned 72 base pairs from the primary cleavage site, adhering to the recommended range of 40-90 nucleotides^19^. Construct assembly was achieved through *BbsI*-mediated golden gate assembly of phosphorylated and annealed oligonucleotide guide sequences (Thermo Fisher), following optimised protocols for PEA1 or pDG459 assembly^20, 21^. Subsequent plasmid purification was performed using the QIAprep Spin Miniprep Kit (Qiagen), and validation of oligonucleotide insertion was carried out through Sanger sequencing (AGRF). Oligonucleotide sequences are listed in **Table 1**.

**Table 1.**
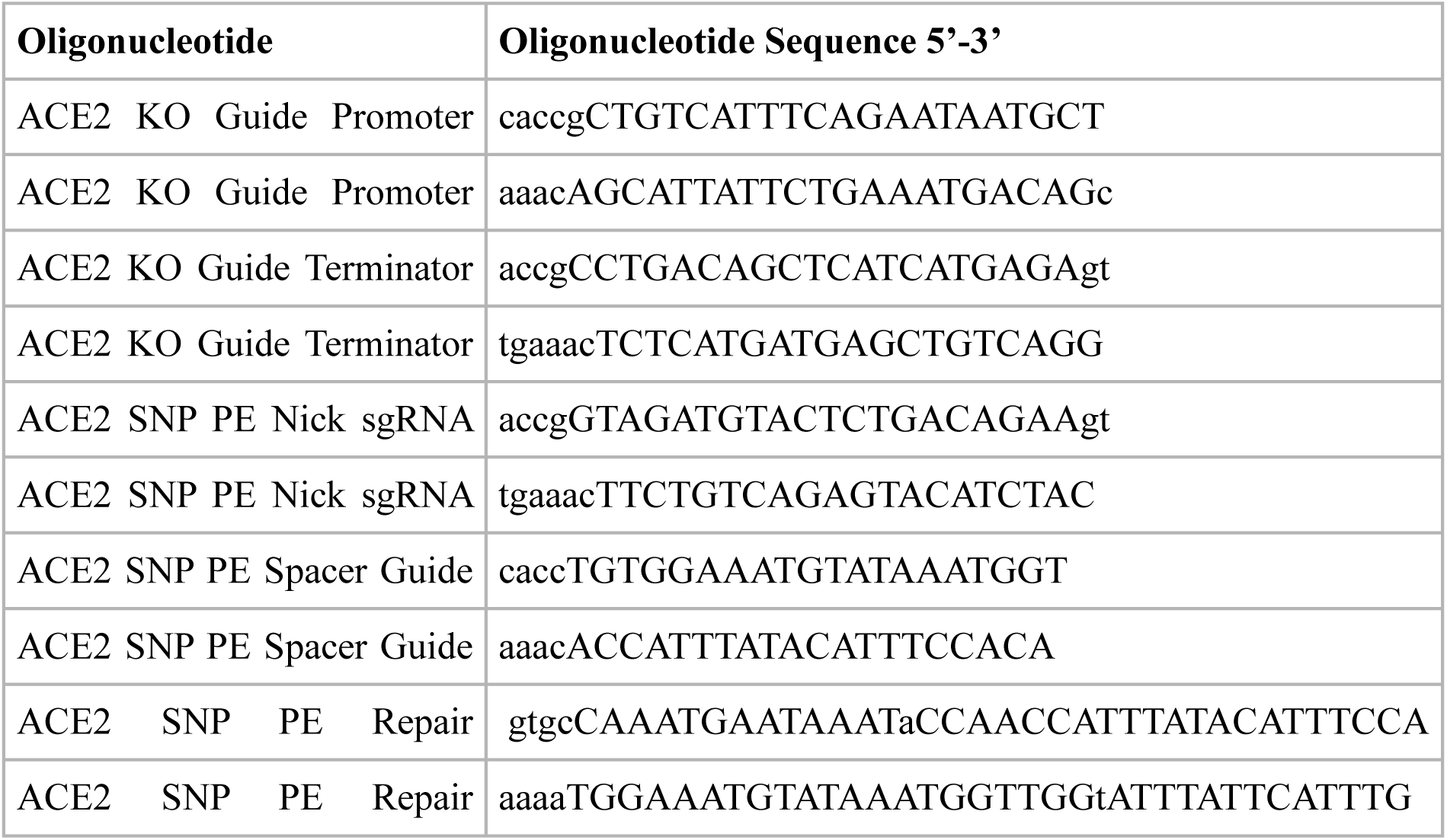
Sequences of oligonucleotides used to generate plasmids for organoid models.

### TSC genetic modification

TSCs were transfected with *ACE2* KO and *ACE2* SNP plasmids using DNAfectin Plus Transfection Reagent (abm), according to manufacturer’s instructions. Briefly, media was aspirated from the TSC culture wells prior to the addition of the DNA-transfection complex. Control wells received only the transfection reagent without DNA plasmid. After 6 h the transfection solution was removed, and complete medium was added. After 24 hours, puromycin (0.2 µg/mL) was added to culture media to begin antibiotic selection of successfully transfected cells. Puromycin selection continued for 14 days of culture (until cells reached confluence).

### Assessment of Time to Confluence and Cell Death

Immediately after culture with puromycin, gene-edited TSCs were passaged using TrypLE Reagent (TrypLE, Gibco), according to manufacturer’s instructions, then replated at a density of ∼20% (1:5 dilution) per well. Cells were then grown in the Incucyte® SX5 for 4 days (until confluence). Cell confluence percentage and cell death were assessed using Incucyte analysis software. Once cells reached confluence, TSCs were removed from the Incucyte and organoids were created.

### Organoid formation

Placental organoids were established as per Haider, *et al.*^18^, with minor modifications. Briefly, gene-edited TSCs were trypsinised (TrypLE, Gibco) and washed in ice-cold advanced Dulbecco’s Modified Eagle Medium (DMEM, Gibco). Cells were resuspended in organoid media: ice-cold advanced DMEM supplemented with 1×B-27, 1×ITS-X, 10 mM HEPES (Gibco), 2 mM glutamine (Gibco), 1 µM A8391, 100 ng/ml rhEGF and 3 µM CHIR99021. Vitrogel (ORGANOID-1, The Well BioScience) was added to reach a 2:1 Vitrogel to organoid media ratio. 75µL of the resultant mixture was plated in a 6-well culture plate, forming a dome in the centre of the well. After incubation (37°C, 2 h), 1.5mL organoid media (37°C) was added to each well. Media was changed every 5 days until organoids reached confluence, at which point they were harvested for RNA and protein experiments.

### Genomic DNA (gDNA) extraction

gDNA extraction protocol was adapted from Ghatak, *et al*.^22^. Prior to organoid generation, a subset of TSCs from each gene-edited patient sample was harvested and gDNA was extracted by incubating cells with a phenol:chloroform:isoamyl alcohol solution (25:24:1). After centrifugation (10 mins, 10000 xg, 4°C), the upper aqueous layer was collected and transferred to a new tube. RNase A (0.1mg; Lucigen) was added. Isopropanol (4°C, 1:1 volume ratio) and sodium acetate (4°C, 3M, 1:10 volume ratio) were added to the mixture prior to precipitation (−20°C, 1 h). The sample was then centrifuged (10000 xg, 10 mins, 4°C) before the supernatant was decanted. The pellet was washed in 250µL 75% EtOH prior to centrifugation (10000 xg, 10 mins, 4°C). The supernatant was again decanted, and the pellet was left to air-dry, then resuspended in nuclease-free H_2_O (30µL).

### Genotyping for *ACE2* rs2074192

Genotyping was conducted by the Australian Genome Research Facility (AGRF) using the Sequenom MassARRAY system, as per Zhou, *et al*.^23^.

### RNA Extraction

Culture media was aspirated from organoid wells and organoids were gently washed with phosphate-buffered saline (PBS). Tissue was disrupted by homogenizing for 3.5 mins at 30 Hz (TissueLyser, QIAGEN) in 600μL Buffer RLT Plus (RNeasy Plus Mini Kit; QIAGEN, Victoria, Australia). Total RNA was extracted from the supernatant using TRIzol reagent as per manufacturer’s instructions and as outlined by Rio, *et al.*^24^. The purity and integrity of extracted RNA samples were determined using the NanoDrop Spectrophotometer (Thermo Fisher Scientific) and samples used had 260:280 and 260:230nm ratios greater than 1.9.

### cDNA Synthesis and Quantitative Polymerase Chain Reaction (qPCR)

As previously described^25^, synthesis of complementary DNA (cDNA) was conducted beginning with 1μg of total RNA using the QuantiNova Reverse Transcription Kit (QIAGEN) according to the manufacturer’s protocol. qPCR was conducted with SYBR Green (QIAGEN) according to manufacturer’s instructions, with YWHAZ (*YWHAZ*) and β-actin (*ACTINB*) as housekeeping genes (for primer sequences, see Table 2). Denaturation was performed at 95°C for 10 secs, annealing at 53°C for 45 secs, and extension at 72°C for 30 secs, for a total of 50 cycles. qPCR results were analysed using the 2^-ΔΔCT^ method^26^.

**Table 2.**
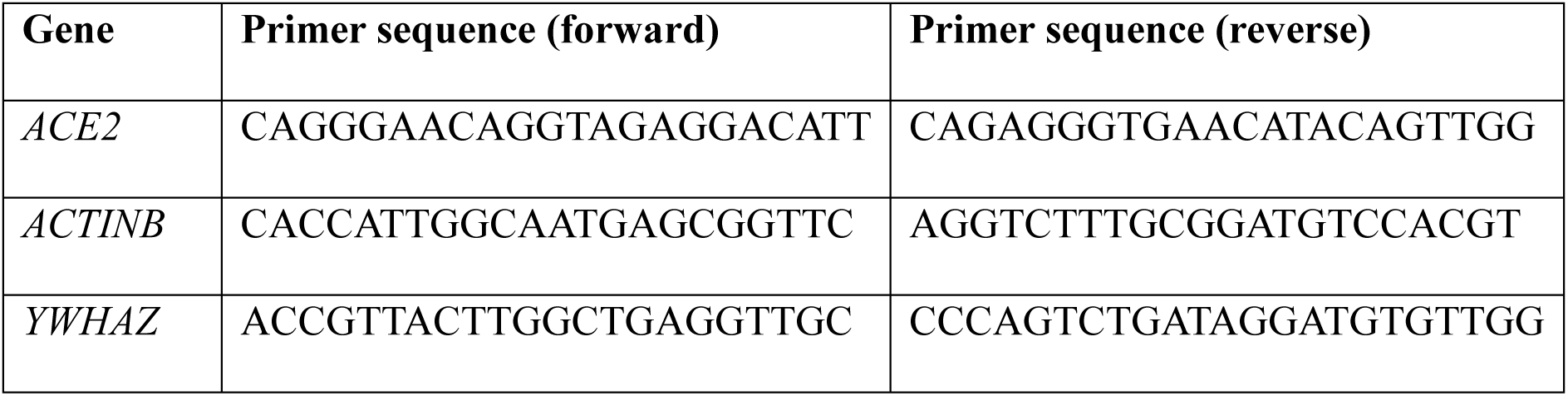
Sequences for qPCR primers.

### Protein extraction

Total protein was extracted from gene-edited placental organoids using a radioimmunoprecipitation assay (RIPA) lysis and extraction buffer. Organoids were harvested from culture wells and each sample was placed in 400µL ice cold RIPA buffer with added Complete Mini Protease Inhibitor Cocktail tablet (Roche Diagnostics Australia) and phenylmethylsulfonyl fluoride (0.4 pmol). Samples were incubated (10 mins, 4°C), vortexed, centrifuged (16000 xg, 10 mins, 4°C) and supernatants collected. Protein concentration was measured via a Bradford Assay using the SpectraMax iD5 (Molecular Devices) at 595 nm absorbance.

### SDS-PAGE

As previously described^25^, samples were denatured and reduced in Laemmli 4x buffer (GTX16355, GeneTex) and 8% 2-mercaptoethanol (5 mins, 95°C). Samples were loaded onto either a 4–20% Mini-PROTEAN^®^ TGX Stain-Free™ Protein Gel (4568094, Bio-Rad) or 4– 20% Mini-PROTEAN^®^ TGX™ Precast Protein Gel (4561094, Bio-Rad), along with a molecular weight marker (GTX49384, GeneTex). SDS-PAGE was performed (1 h, 100 V) in 1x Running buffer (pH 8.3) using a Mini-PROTEAN Tetra Vertical Electrophoresis Cell (Bio-Rad). Stain-Free Protein gels were checked for complete protein separation using the ChemiDoc Touch Imaging System (Bio-Rad).

### Transfer

Protein was transferred to a PVDF membrane over 16 h (27V, 4°C) in transfer buffer (25 mM Trizma Base, 190 mM glycine, 20% methanol; pH 8.3) using the Criterion™ blotter (Bio-Rad). Successful transfer of protein was checked by Ponceau S staining of the membrane, as well as either Coomassie Blue staining (for the TGX Precast Protein Gel) or the ChemiDoc Touch Imaging System (Bio-Rad) (for the TGX Stain-Free™ Protein Gel).

### Blotting

Membrane was blocked in Tris Buffered Saline Tween20 (TBST) with 5% skim milk (2 h, 25°C). Membrane was incubated in blocking buffer (1 h, 25°C) with Anti-ACE2 Polyclonal Antibody (ab15348, Abcam) at 1:300 dilution, then washed 3-4 times with TBST, and incubated with a goat anti-rabbit Immunoglobulin/HRP secondary antibody (P044801-2, Dako) at 1:5000 in the blocking buffer. Membrane was washed 3-4 times in TBST, then incubated in Clarity Western ECL Substrate (1705060, Bio-Rad) for 5 mins and imaged. The density of each band (determined by the ChemiDoc Touch Imaging System) was determined prior to analysis using ImageLab software from Bio-Rad™. Analysis controlled protein loading for each sample by normalizing using stain-free total protein quantification^27^ and was further normalized to an internal control sample (pooled TSCs) on each membrane. Samples were run in duplicate and averaged for final analysis.

### Enzyme-Linked Immunosorbent Assay (ELISA)

A commercially available ELISA was used to measure the concentration of ACE (DY929, Duoset R&D Systems) according to manufacturer’s instructions, using methods previously described^28^.

### ACE2 activity assay

ACE2 enzyme activity was assessed according to Xiao and Burns^29^ and Tamanna, *et al.^1^*. Briefly, samples (15µL; cell lysate or cell media) were diluted with enzyme buffer (1M NaCl, 75mM Tris-HCl, 5mM ZnCl_2_; pH 6.5) with protease inhibitors (10µM captopril, 5µM amastatin, 10µM bestatin; Sigma Aldrich, 10µM Z-prolyl-prolinal; Enzo Life Sciences). MCA-Ala-Pro-Lys(Dnp)-OH (AnaSpec, #AS-60757), an ACE-2 specific fluorescent substrate, was diluted in enzyme buffer and added to samples (final concentration 50µM). Samples were run in duplicate along with a blank control in a 96-well black microplate. The plate was covered and incubated on a plate shaker (24 h, 25°C).

The ACE2 standard curve was generated using human recombinant ACE2 protein (R&D Systems, #933-ZN). For each concentration of ACE2 standard, two wells were dedicated to measuring total ACE2 activity and two dedicated to measuring activity in the presence of DX600 (AnaSpec, #AS-62337), an ACE2 inhibitor. Relative Fluorescence Units (RFU) measured the product MCA-Ala-Pro over 24 h from the substrate MCA-Ala-Pro-Lys(Dnp)-OH as cleaved by ACE2. Fluorescence readings (CLARIOstar Plus, BMG LabTech) were taken with an excitation wavelength of 320nM, emission wavelength of 405nM.

ACE2 activity was calculated by comparing the RFU of known concentrations of ACE2 standard to the RFU of our samples. As such, ACE2 activity is reported as activity/ng/mL of ACE2.

### Measurement of organoid diameter and assessment of growth

Images of organoid wells were captured using Brightfield microscopy (Olympus Life Science). All wells were imaged in the same orientation. Diameters were measured using QPath Software horizontally across each organoid for n=100 organoids per group, per patient. Symmetrical growth was assessed by comparing the horizontal diameter for each organoid with the vertical diameter for each organoid, for n=100 organoids per group, per patient.

### Statistical Analysis

Statistical analysis for differences between groups was undertaken using SPSS Statistics Software. Outliers were removed from the data using a Grubbs’ test. Data were assessed for normal distribution and a two-way ANOVA test was conducted. Differences between groups were considered significant for *p*<0.05. All tests of statistical significance are two-sided, and adjustments were made for multiple comparisons.

## Results

### *ACE2* knockout (KO) placental organoids

*ACE2^+/-^ and ACE2*^-/-^ (hereafter referred to as ‘*ACE2* KO’) placental organoids were successfully generated. *ACE2^+/+^ and ACE2*^+/-^ organoids expressed significantly more *ACE2* mRNA than *ACE2* KO organoids (*p*=0.00155, *p*=0.0324 respectively; **Figure 3A**). No significant difference in *ACE2* mRNA abundance was detected between *ACE2^+/+^ and ACE2*^+/-^ organoids. ACE2 protein was not detected in cell media nor cell lysate of *ACE2* KO organoids (nil detected). Therefore, ACE2 protein was significantly increased in cell media (**Figure 3B**) and cell lysate (**Figure 3C, 3D**) of *ACE2^+/+^ and ACE2*^+/-^ organoids compared to *ACE2* KO organoids (*p*<0.0001 for all). No significant difference in ACE2 protein abundance was detected between *ACE2^+/+^ and ACE2*^+/-^ organoids in cell media or cell lysate.

**Figure 3.**
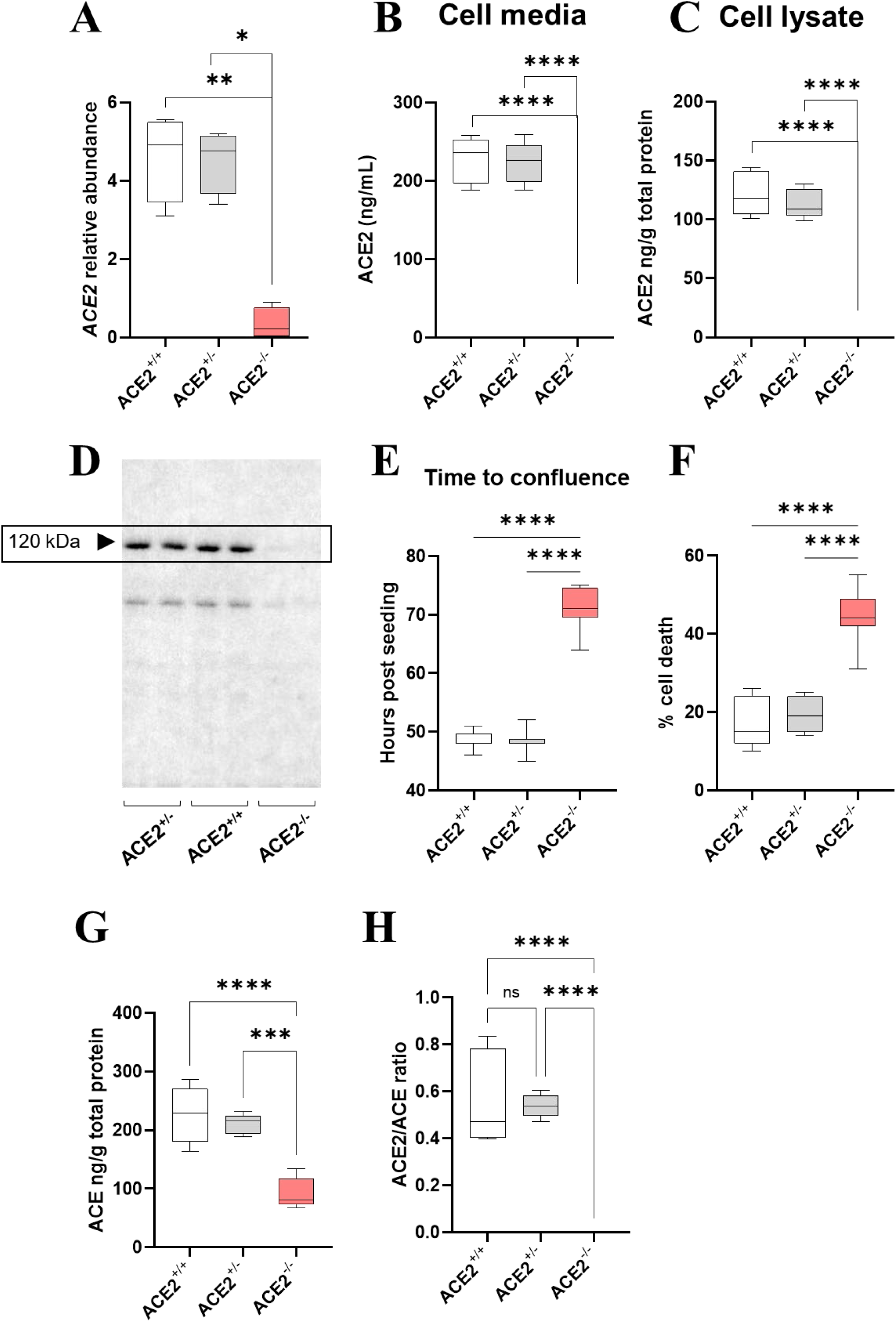
The abundance of **(A)** ACE2 mRNA, **(B)** ACE2 protein in cell media and **(C)** ACE2 protein in cell lysate of ACE^+/+^, ACE^+/-^ and ACE2^-/-^ (ACE2 KO) organoids. **(D)** Representative WB showing ACE2 protein expression in cell lysate of ACE^+/+^, ACE^+/-^ and ACE2 KO organoids. **(E)** The time to confluence, in hours post seeding, and **(F)** percentage cell death in ACE^+/+^, ACE^+/-^ and ACE2 KO trophoblast stem cells (TSCs). **(G)** Level of ACE protein in cell lysate and **(H)** ACE2/ACE (ACE2:ACE) ratio (from cell lysate levels) of ACE^+/+^, ACE^+/-^ and ACE2 KO organoids. Data are presented as a 10-90 percentile interleaved box-and-whisker plot. White bars denote ACE^+/+^ group, grey bars denote ACE^+/-^ group, red bars denote ACE2 KO group. ns indicates non-significance, *p<0.05, **p<0.01, ***p<0.001, ****p<0.0001. (n=9)

ACE2 knockout leads to slower cell growth and cell death in placental organoids *ACE2^+/+^* and *ACE2^+/-^* TSCs reached confluence at an average of 48.5 and 48.3 hours post seeding, respectively. In contrast, *ACE2* KO TSCs reached confluence much later at an average of 71.1 hours post seeding. *ACE2^+/+^* and *ACE2^+/-^* TSCs reached confluence significantly faster than *ACE2* KO TSCs (*p*<0.0001 for all, **Figure 3E**).

*ACE2^+/+^* and *ACE2^+/-^* TSCs had average cell death proportions of 17% and 19%, respectively. In contrast, *ACE2* KO TSCs had an average cell death proportion of 44%. *ACE2^+/+^* and *ACE2^+/-^* TSCs had significantly lower proportions of cell death than *ACE2* KO TSCs (*p*<0.0001 for all, **Figure 3F**).

### *ACE2* KO markedly reduces ACE protein expression in placental organoids

*ACE2^+/+^* and *ACE2^+/-^* organoids expressed significantly more ACE protein than *ACE2* KO organoids (*p*<0.0001, *p*=0.0003, respectively; **Figure 3G**). Given the absence of ACE2 protein in *ACE2* KO organoids, the ACE2:ACE ratio was significantly higher in *ACE2^+/+^* and *ACE2^+/-^* organoids compared to *ACE2* KO organoids (*p*<0.0001 for both; **Figure 3H**) but remained unchanged between *ACE2^+/+^* and *ACE2^+/-^* organoids (no significant difference).

*ACE2* KO reduces diameter and induces asymmetrical growth in placental organoids Placental organoids tend to exhibit a spherical structure^17, 18^. However, with the *ACE2* knockout we observed significantly smaller placental organoids with ‘squashed’, i.e. asymmetrical, structures (**Figures 4A-C**). We have therefore measured the horizontal and vertical diameters of organoids from each genotype to determine the effect of *ACE2* on growth and symmetry.

**Figure 4.**
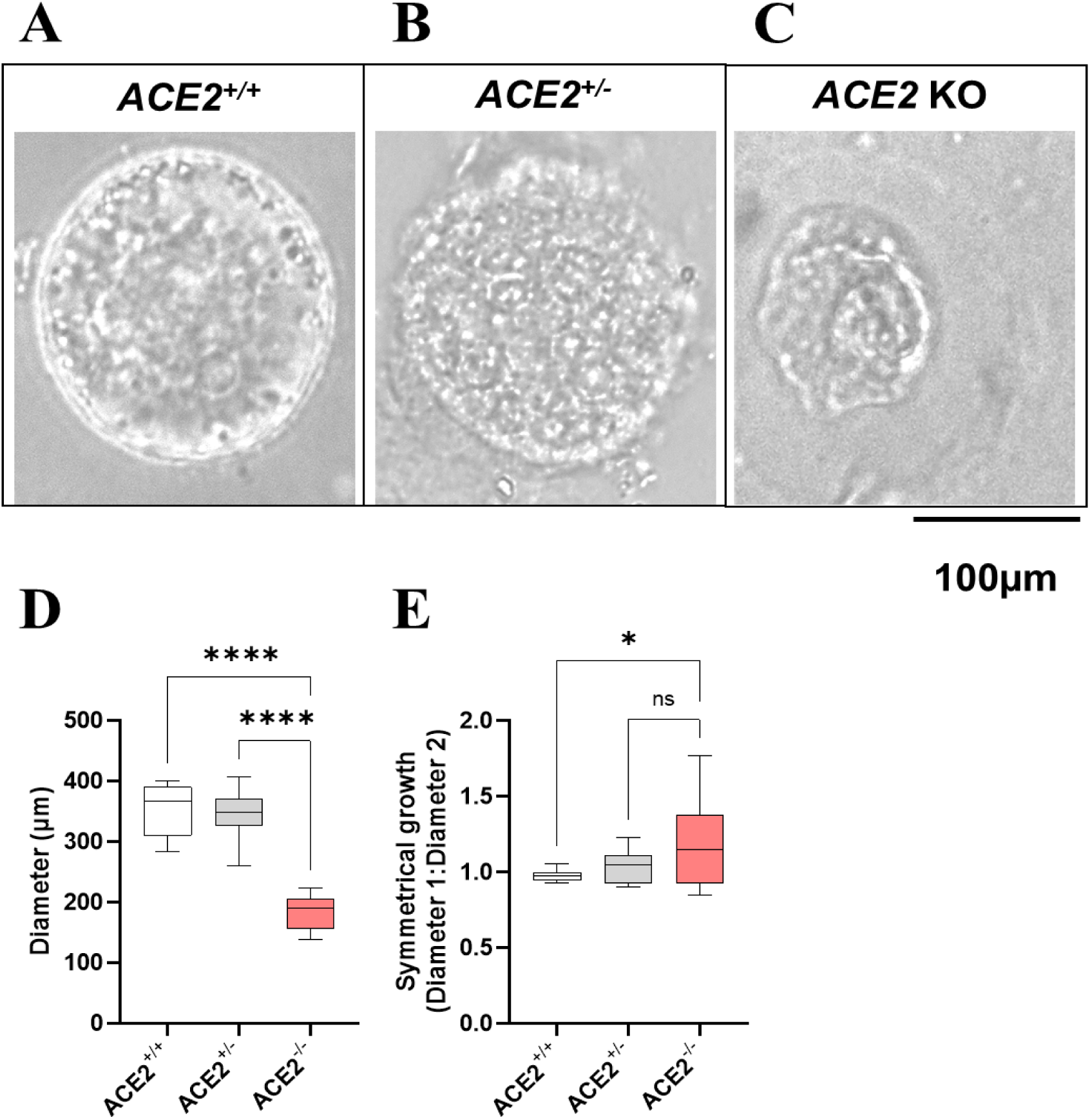
Representative image (Brightfield) of symmetrical organoid growth (**A**) ACE2^+/+^, (**B**) ACE2^+/-^ and (**C**) asymmetrical organoid growth (ACE2 KO). **(D)** The average diameter (µm) of ACE^+/+^, ACE^+/-^ and ACE2 KO organoids. **(E)** The ratio of diameter 1 (horizontal) to diameter 2 (vertical) of ACE^+/+^, ACE^+/-^ and ACE2 KO organoids as a measure of symmetrical organoid growth. A value of 1.0 indicates perfect symmetry. Data are presented as a 10-90 percentile interleaved box-and-whisker plot. White bars denote ACE^+/+^ group, grey bars denote ACE^+/-^ group, red bars denote ACE2 KO group. ns indicates non-significance, *p<0.05, ****p<0.0001. (n=9)

*ACE2^+/+^*and *ACE2^+/-^*organoids had significantly larger diameters compared to *ACE2* KO organoids (*p*<0.0001 for all, **Figure 4D**).

Organoids were significantly less symmetrical in *ACE2* KO organoids compared with *ACE2^+/+^* organoids (*p*=0.0186, **Figure 4E**) only.

### Placental organoids with *ACE2* rs2074192 induced SNP

TSCs were successfully gene-edited to induce the *ACE2* rs2074192 SNP. The plasmid accuracy was confirmed by Sanger sequencing (**Figure 5**), which verified a single base substitution from C to T at position 15564667 on the X chromosome (reference GRCh38.p2 38.2/144), *ACE2* gene. Transfected trophoblast stem cells were genotyped (see methods) to verify successful SNP candidates, from which organoids were created. Three genotype groups were created: TT, heterozygous SNP (CT) and homozygous SNP (TT).

**Figure 5.**
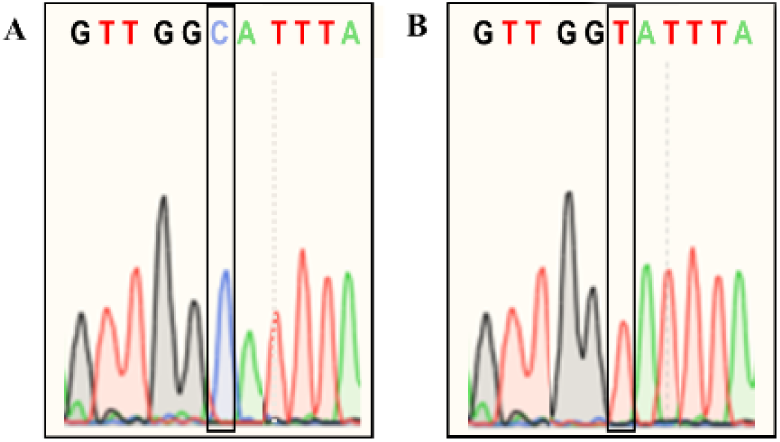
Successful gene-editing at position 6: the reference C allele (A) and the alternate T allele (B).

### *ACE2* rs2074192 TT organoids, but not rs2074192 CT organoids, have increased *ACE2* mRNA and ACE2 protein

TT organoids had significantly higher expression of *ACE2* mRNA compared with both CC and CT organoids (*p*=0.0040, *p*=0.0141, respectively; **Figure 6A**). TT organoids also had significantly higher levels of ACE2 protein than CC and CT organoids in both the cell media (*p*=0.0013, *p*=0.0169, respectively; **Figure 6B**) and cell lysate (*p*=0.0018, *p*=0.0040, respectively; **Figure 6C, 6D**). There was no significant difference in abundance of *ACE2* mRNA, nor ACE2 protein in cell media or lysate, between CC and CT organoids.

**Figure 6.**
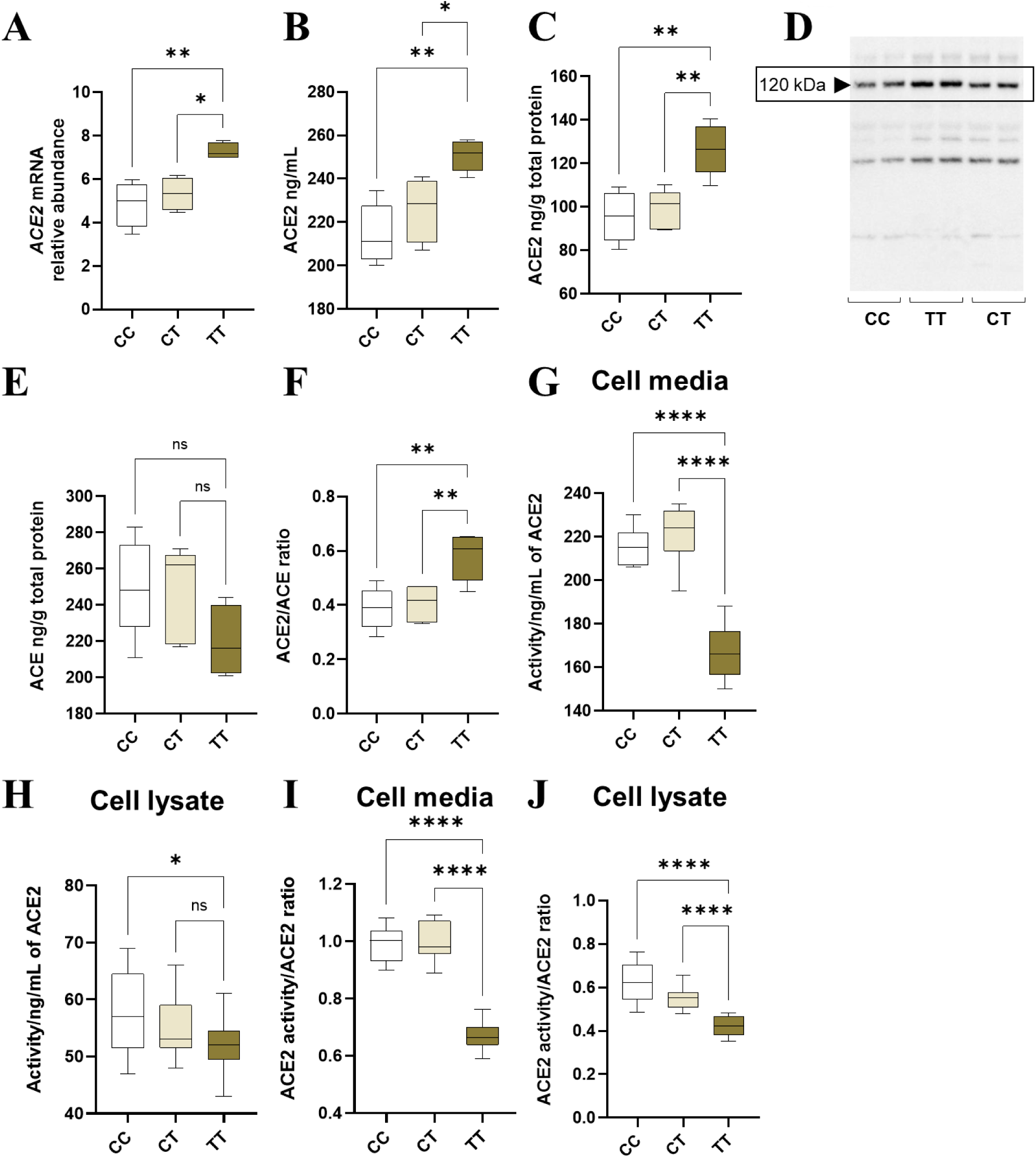
The abundance of **(A)** ACE2 mRNA, **(B)** ACE2 protein in cell media and **(C)** ACE2 protein in cell lysate; **(D)** Representative WB showing ACE2 protein expression in cell lysate of CC, TT and CT organoids. **(E)** Level of ACE protein in cell lysate, **(F)** ACE2/ACE ratio (from cell lysate levels); Enzyme activity of ACE2 in **(G)** cell media and (**H**) cell lysate, and ratio between ACE2 activity and ACE2 levels in (**I**) cell media and (**J**) cell lysate of CC, CT and TT organoids. Data are presented as a 10-90 percentile interleaved box-and-whisker plot. White bars denote CC group, beige bars denote CT group, brown bars denote TT group. ns indicates non-significance, *p<0.05, **p<0.01, ***p<0.001, ****p<0.0001. (n=9)

Time to confluence, and percentage cell death, were also assessed, however no significant differences were detected between groups (data not shown).

### *ACE2* rs2074192 genotype did not change ACE protein expression, but increased the ACE2/ACE ratio in TT organoids

There was no significant difference in ACE protein level (measured in cell lysate) between CC, CT and TT organoids (**Figure 6E**). However, the ACE2/ACE ratio was significantly higher in TT organoids compared to both CC and CT organoids (*p*=0.0049, *p*=0.0093, respectively; **Figure 6F**). There was no significant difference in ACE protein expression, nor ACE2/ACE ratio, between CC and CT organoids.

### ACE2 activity is decreased in *ACE2* rs2074192 TT organoids

ACE2 enzyme activity was significantly reduced in the cell media of TT organoids compared to both CC and CT organoids (*p*<0.0001 for both; **Figure 6G**). There was no significant difference in ACE2 activity in the cell media of CT organoids compared to CC organoids. ACE2 enzyme activity was significantly reduced in the cell lysate of TT organoids compared to CC organoids (*p*=0.0303; **Figure 6H**). There was no significant difference in ACE2 activity in the cell lysate of CT organoids compared to TT organoids, nor compared to CC organoids.

The ratio between ACE2 activity and ACE2 levels was significantly decreased in the cell media of TT organoids compared to both CC and CT organoids (*p*<0.0001 for both; **Figure 6I**), while there was no significant difference between CT and CC organoids. The ratio between ACE2 activity and ACE2 levels was significantly decreased in the cell lysate of TT organoids compared to both CC and CT organoids (*p*<0.0001 for both; **Figure 6J**), while there was no significant difference in the ratio between CT and CC organoids.

### *ACE2* rs2074192 TT organoids have reduced diameters and asymmetrical growth

The diameter of TT organoids was significantly reduced compared to CC organoids (*p*=0.0141; **Figure 7A**) only. Organoid growth was significantly less symmetrical in TT organoids compared to both CC and CT organoids (*p*=0.0077, *p*=0.0415, respectively; **Figure 7B**). There was no significant difference in diameter or symmetrical growth measurements between CC and CT organoids.

**Figure 7.**
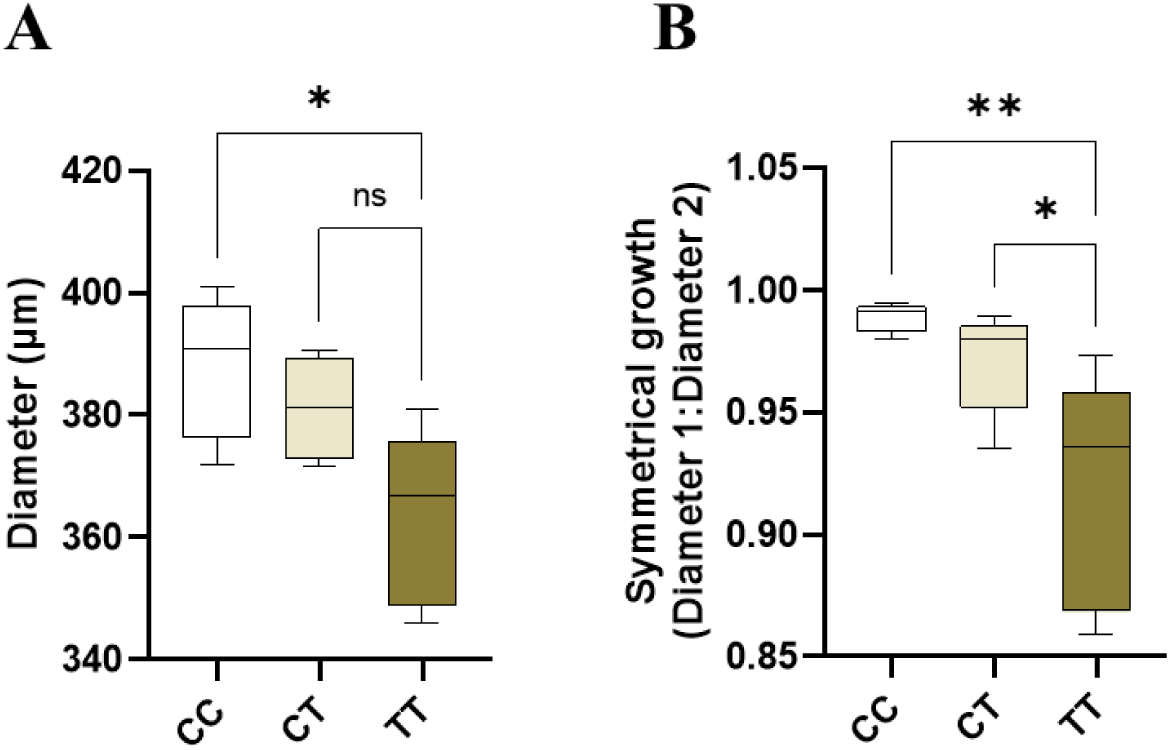
**(A)** The average diameter (µm) of CC, CT and TT organoids. **(B)** The ratio of diameter 1 (horizontal) to diameter 2 (vertical) of CC, CT and TT organoids as a measure of symmetrical organoid growth. A value of 1.0 indicates perfect symmetry. Data are presented as a 10-90 percentile interleaved box-and-whisker plot. White bars denote CC group, beige bars denote CT group, brown bars denote TT group. ns indicates non-significance, *\*p<0.05, **p<0.01. (n=9)*

## Discussion

This study is the first to successfully use gene-editing to stably produce *ACE2* knockout placental organoids, and the first to induce a SNP in placental organoids. Using these models, the study demonstrates that *ACE2* is important for cell growth (as evidenced by changes in the time to confluence and percentage cell death of TSCs), as well as placental organoid size and growth (as evidenced by changes in organoid diameters and symmetry growth measurements). Furthermore, we show that ACE2:ACE ratio is disrupted in both organoid models, resulting in impaired growth and development. Our data are consistent with our hypothesis that *ACE2* plays an important role in placental development, and that ACE2 and ACE need to exist in a finely tuned balance to ensure proper growth and morphogenesis. We have also confirmed that the *ACE2* rs2074192 TT genotype results in increased ACE2 mRNA and protein, and altered ACE2:ACE ratios, which likely mediate the effects on TSC and organoids growth and cell survival.

*ACE2* knockout mouse models of pregnancy have been useful in reinforcing the importance of ACE2 in placentation and pregnancy health. This study has extended our understanding of the role of ACE2 in the placenta, creating the first *ACE2* knockout in primary human placenta cells. Furthermore, we have explored the molecular implications of carriage of a common *ACE2* SNP, rs2074192. *ACE2* variants are associated with pregnancy complications, as well as with a number of other diseases characterised by renin-angiotensin system dysfunction, including COVID-19^13, 14^, hypertension^12, 30^ and heart failure^15^. *ACE2* rs2074192 is a common SNP, with an estimated frequency of C (reference allele) = 0.63 and T (alternate allele) = 0.37, according to a study of 102,615 people (including varied ethnicities) in the Genome Aggregation Database^16^. This suggests that at least one copy of this SNP naturally occurs in over one third of the population, and a recent study indicated that the incidence of the TT genotype in their study population was 19.8%^10^. This highlights the importance of understanding the mechanism by which carriage of the alternate allele increases the risk of disease.

*In silico* analysis of *ACE2* rs2074192 suggests that it increases splicing donor sites, creating a changed secondary RNA structure, affecting protein production and activity. As *ACE2* rs2074192 is an intronic polymorphism, it is likely that it indirectly positively regulates gene expression by altering ACE2 splicing and transcription enhancement^13^. This study confirms that carriage of the TT genotype, but not the CT genotype, results in increased placental ACE2 mRNA and protein expression. This has major implications for the pathogenesis of diseases dependent on ACE2 expression, including pregnancy complications such as preeclampsia^1^ and SGA^1^, and infection with SARS-CoV-2 virus^31^, which can directly infect syncytiotrophoblasts through earlier endometrial cell infection^32^. However, the mechanism of action by which this SNP contributes to pathogenesis requires further study. Both the *ACE2* rs2074192 (C to T) SNP and haplotypes CAGC and TAGT are associated with altered levels of the Ang 1-7 peptide, a direct product of ACE2 activity^10^. As this study did not quantify Ang 1-7 expression, further studies should investigate whether Ang 1-7 is altered as a result of *ACE2* rs2074192 genotype, providing additional information on a potential mechanism for pathogenesis of diseases dependent on ACE2 in T allele carriers.

This study observed that *ACE2* knockout TSCs took longer to reach confluence and had higher percentages of cell death in culture. This finding has implications for pregnancy complications with origins in early gestation, when TSCs play an important role in placental development that underpins subsequent function. During early gestation, the placenta undergoes rapid trophoblast proliferation and differentiation^33^. Not only did this study observe slower TSC growth and higher levels of cell death in *ACE2* knockouts, but *ACE2* knockout organoids had reduced diameters, as well as asymmetrical growth, reflecting impaired organoid development. Interestingly, although no change in TSC growth or cell death was observed with *ACE2* rs2074192 genotype, TT organoids also had smaller diameters and asymmetrical growth.

This impaired growth has concerning implications for placental development. Human placental organoids provide an innovative model for studying placental development *in vitro*. Ethical limitations to study of early-to mid-gestation placentae, created the need for physiologically relevant human placental models with 3D architecture. Current models are limited^34,35,36^ while mouse placenta differs both in structure and endocrine function^37^. Organoids recapitulate all first-trimester placental functions, can differentiate into the expected cell lineages, and are able to be cultured long-term without the loss of these features^38^. For this reason, human placental organoids are the closest possible parallel to studying the early gestation human placenta *in vivo*. *ACE2* gene-editing of placental organoids in this study provides insight into the detrimental effect of *ACE2* knockout and rs2074192 TT genotype on placental development, likely leading to aberrant placentation in human pregnancy.

Whilst an absence of ACE2 is clearly detrimental to cell growth and increases cell death, a complete knockout is highly unlikely in nature and thus is less biologically relevant for causal inference. Therefore, we have compared the knockout and induced SNP models to make inferences most biologically relevant to human pregnancy. Surprisingly, both the *ACE2* KO organoids and the induced SNP organoids exhibited diminished growth and symmetry, despite *ACE2* KO organoids having markedly reduced ACE2 protein, and induced SNP organoids having markedly increased protein, compared to their respective controls. This result suggests that the absolute expression value of ACE2 is the not sole determinant of organoid growth.

One common feature between *ACE2* knockout organoids and TT organoids is the altered ACE2:ACE ratio. Our data suggest that ACE2 and ACE, which are known to partly control cell growth^3^, have more contributions to placental organoid development (as evidenced by organoid symmetry and growth) than simply by affecting cell proliferation and death. This is evident from our observation of unchanged TSC growth and cell death levels in TT TSCs, despite them also having altered ACE2 and ACE expression.

The mechanism for this unexpected increase in ACE2 protein, but decreased organoid growth and symmetry, could be due to the reduced ACE2 enzyme activity observed in TT organoids. *ACE2* rs2074192 is an intronic SNP, which does not directly impact protein coding regions of the genome. Therefore, the contribution of this SNP to ACE2 protein increase is indirect, likely through a complex mechanism not yet elucidated. We hypothesise that the excess of ACE2 protein in TT organoids has resulted in a compensatory response in the cell, reducing the *activity* of ACE2 in an attempt to re-establish an ACE2:ACE equilibrium. This theory is supported by the literature, showing that *ACE2* rs2074192 is associated with increased risk of SGA^5^, a condition in which inadequate placentation often leads to smaller placentae and a smaller fetus. Furthermore, in SGA, ACE2 activity is significantly reduced^1^.

This study indicates that, whilst ACE2 expression is significant, the activity of the enzyme and maintenance of the ACE2 and ACE in equilibrium (in this paper termed an ACE2:ACE ratio) may be of greater importance. Studies investigating pregnancy complications support this conclusion. In preeclampsia, a potentially life-threatening complication characterised by maternal hypertension and multi-organ failure, ACE2 activity reflected by the ACE2:ACE ratio is higher than normal, impairing placental development and function^1^. In SGA, placental expression of ACE and ACE2 at delivery are both higher compared with uncomplicated pregnancy. Birthweight centiles are negatively correlated with maternal plasma ACE and ACE2 levels in mid-pregnancy, suggesting that tight regulation of these enzymes is critical to ensure pregnancy health^1^. As previously mentioned, the ratio of ACE2 activity to ACE2 levels in maternal plasma is significantly reduced in SGA, indicating that ACE2 operates with reduced efficiency in this condition^1^. This could be due to the increased presence of an endogenous inhibitor^39^ in SGA, although this has never been investigated. Placental ACE2 deficiency, and the associated elevation in placental Ang II levels, negatively impact pregnancy by impairing maternal weight gain and restricting fetal growth^2^. This implies the balance of ACE2:ACE supersedes the importance of ACE2 levels in isolation in determining fetal growth. Indeed, herein we observed similar reductions in organoid diameter and growth between the complete *ACE2* knockout and rs2074192 models, despite the difference in absolute ACE2 expression values.

Interestingly, this study showed that *ACE2* rs2074192 TT organoids not only had elevated ACE2 protein within cells (as assessed by cell lysate protein levels) but also secreted more ACE2 protein into the cell media. Whilst ACE2 is typically a membrane-bound peptidase, it can also be shed from the membrane^40^, producing soluble ACE2 (sACE2). It should be noted that this study utilised only an ACE2 antibody specific to the membrane-bound ACE2 protein and we cannot comment on the production of sACE2 by these organoid models. Future studies will utilise a specific sACE2 antibody to examine this.

A limitation of this study is that it only utilised placental tissue from female fetuses. This is because placental expression of ACE and ACE2 is fetal sex dependent. Both ACE and ACE2 are more highly expressed in placentae from female compared to male fetuses^41^. For ACE2, this is likely due to the fact that *ACE2*, located on the X-chromosome, escapes X chromosome inactivation, thus often two copies are expressed instead of one in females^42^. The balance of downstream effector peptides Ang II (via ACE) and Ang 1-7 (via ACE2) is also influenced by fetal sex. In early gestation, Ang 1-7:Ang II ratios are reduced in placenta from females, indicating less Ang 1-7 production as a result of reduced ACE2 activity^43^. Female babies are more likely to be growth restricted than males^44, 45^. In this study, the organoid growth symmetry only significantly changed between the *ACE2^+/+^* and total knockout, but not *ACE2^+/-^* and the knockout. This could indicate that in males, where only one *ACE2* gene is present (due to the presence of only one X chromosome), there may be differences in placental development. Thus, *ACE2* fetal sex-specific molecular roles require further investigation. Future studies should include male samples, as well as females to expand on the findings from this study.

Our data clearly demonstrate that ACE2 plays an important role in placental organoid development, and its expression is increased with the rs2074192 TT genotype. Importantly, ACE2 enzyme activity and the ACE2:ACE ratio appeared to make more difference to the growth and development of TSCs and placental organoids than the absolute expression values of ACE2 alone. We anticipate that this study will be used to facilitate a deeper understanding of the role of ACE2 in placental development. The gene-editing technique in placental organoids that we have developed could be used to study the roles of many other genes and their variants on placental growth and function.

## Acknowledgments

We thank the women who consented to their placental tissue being used for this research. We thank Imre Schene, of UMC Utrecht, for his advice and expertise in developing genetically edited organoids. Figures were made using Biorender.com.

## Author contributions

A.L.A conceptualised this work, performed experimental work and formal analysis of data, original draft preparation, and review and editing. B.D. and M.K. provided valuable contributions to methodology optimisation, and reviewed and edited the manuscript. C.J.L., P.Q.T. and F.A. designed and generated plasmids used for gene-editing, and reviewed and edited the manuscript. T.J-K., M.D.S. and K.G.P. reviewed and edited the manuscript. C.T.R. provided supervision, funding acquisition, interpretation of data, review and editing of the manuscript. All authors contributed to the article and approved the submitted version.

## References

1. Tamanna S, Clifton VL, Rae K, van Helden DF, Lumbers ER, Pringle KG. Angiotensin converting enzyme 2 (ACE2) in pregnancy: preeclampsia and small for gestational age. Frontiers in physiology, 1259 (2020).

2. Bharadwaj MS, et al. Angiotensin-converting enzyme 2 deficiency is associated with impaired gestational weight gain and fetal growth restriction. Hypertension 58, 852–858 (2011).

3. Goyal N, Yellon SM, Longo LD, Mata-Greenwood E. Placental gene expression in a rat ‘model’ of placental insufficiency. Placenta 31, 568–575 (2010).

4. Yamaleyeva LM, et al. Uterine artery dysfunction in pregnant ACE2 knockout mice is associated with placental hypoxia and reduced umbilical blood flow velocity. American Journal of Physiology-Endocrinology and Metabolism 309, E84–E94 (2015).

5. He J, et al. Fetal but not maternal angiotensin converting enzyme (ACE)-2 gene Rs2074192 polymorphism is associated with increased risk of being a small for gestational age (SGA) newborn. Kidney and Blood Pressure Research 43, 1596–1606 (2018).

6. Delforce SJ, Lumbers ER, Ellery SJ, Murthi P, Pringle KG. Dysregulation of the placental renin– angiotensin system in human fetal growth restriction. Reproduction 158, 237–245 (2019).

7. Neves LA, et al. ACE2 and ANG-(1-7) in the rat uterus during early and late gestation. American Journal of Physiology-Regulatory, Integrative and Comparative Physiology 294, R151–R161 (2008).

8. Song W, Wang H, Ma L, Chen Y. Associations between the TNMD rs4828038 and ACE2 rs879922 polymorphisms and preeclampsia susceptibility: a case-control study. Journal of Obstetrics and Gynaecology, 1–5 (2022).

9. Rice G, Jones A, Grant P, Carter A, Turner A, Hooper N. Circulating activities of angiotensin-converting enzyme, its homolog, angiotensinconverting enzyme 2, and neprilysin in a family study. Hypertension 48, 914–992 (2006).

10. Chen Y, et al. Relationship between genetic variants of ACE 2 gene and circulating levels of ACE 2 and its metabolites. Journal of Clinical Pharmacy and Therapeutics 43, 189–195 (2018).

11. Valdes G, et al. Distribution of angiotensin-(1-7) and ACE2 in human placentas of normal and pathological pregnancies. Placenta 27, (2006).

12. Luo Y, et al. Association of ACE2 genetic polymorphisms with hypertension-related target organ damages in south Xinjiang. Hypertension Research 42, 681–689 (2019).

13. Pouladi N, Abdolahi S. Investigating the ACE2 polymorphisms in COVID-19 susceptibility: An in silico analysis. Molecular Genetics & Genomic Medicine 9, e1672 (2021).

14. Sabater Molina M, et al. Polymorphisms in ACE, ACE2, AGTR1 genes and severity of COVID-19 disease. PloS one 17, e0263140 (2022).

15. Patel SK, et al. Association of ACE2 genetic variants with blood pressure, left ventricular mass, and cardiac function in Caucasians with type 2 diabetes. American journal of hypertension 25, 216–222 (2012).

16. Karczewski KJ, et al. The mutational constraint spectrum quantified from variation in 141,456 humans. Nature 581, 434–443 (2020).

17. Dietrich B, et al. NOTCH3 signalling controls human trophoblast stem cell expansion and differentiation. Development 150, dev202152 (2023).

18. Haider S, et al. Self-renewing trophoblast organoids recapitulate the developmental program of the early human placenta. Stem cell reports 11, 537–551 (2018).

19. Anzalone AV, et al. Search-and-replace genome editing without double-strand breaks or donor DNA. Nature 576, 149–157 (2019).

20. Adikusuma F, Pfitzner C, Thomas PQ. Versatile single-step-assembly CRISPR/Cas9 vectors for dual gRNA expression. PLOS ONE 12, e0187236 (2017).

21. Adikusuma F, et al. Optimized nickase-and nuclease-based prime editing in human and mouse cells. Nucleic acids research 49, 10785–10795 (2021).

22. Ghatak S, Muthukumaran RB, Nachimuthu SK. A simple method of genomic DNA extraction from human samples for PCR-RFLP analysis. Journal of biomolecular techniques: JBT 24, 224 (2013).

23. Zhou A, et al. The association of maternal ACE A11860G with small for gestational age babies is modulated by the environment and by fetal sex: a multicentre prospective case–control study. Molecular Human Reproduction 19, 618–627 (2013).

24. Rio DC, Ares M, Hannon GJ, Nilsen TW. Purification of RNA using TRIzol (TRI reagent). Cold Spring Harbor Protocols 2010, pdb. prot5439 (2010).

25. Arthurs AL, et al. Placental Inflammasome mRNA Levels Differ by Mode of Delivery and Fetal Sex. Frontiers in Immunology 13, (2022).

26. Livak KJ, Schmittgen TD. Analysis of relative gene expression data using real-time quantitative PCR and the 2− ΔΔCT method. methods 25, 402–408 (2001).

27. Hammond M, Kohn J, Oh K, Piatti P, Liu N. A method for greater reliability in Western blot loading controls—Stain-Free total protein quantitation. Bio-Rad Bulletin 6360, (2013).

28. Arthurs A, Lumbers ER, Delforce SJ, Mathe A, Morris BJ, Pringle KG. The role of oxygen in the regulation of miRNAs in control of the placental renin-angiotensin system. Molecular Human Reproduction 25, 206–217 (2019).

29. Xiao F, Burns KD. Measurement of angiotensin converting enzyme 2 activity in biological fluid (ACE2). In: Hypertension). Springer (2017).

30. Patel S, Velkoska E, Freeman M, Wai B, Lancefield T, Burrell L. From gene to protein-experimental and clinical studies of ACE2 in blood pressure control and arterial hypertension. Front Physiol 5, 227 (2014).

31. Zhou P, et al. A pneumonia outbreak associated with a new coronavirus of probable bat origin. Nature 579, 270–273 (2020).

32. Chen J, et al. A placental model of SARS-CoV-2 infection reveals ACE2-dependent susceptibility and differentiation impairment in syncytiotrophoblasts. Nature Cell Biology 25, 1223–1234 (2023).

33. Gude N, Roberts CT, Kalionis B, King RG. Growth and function of the normal human placenta. Thrombosis Research 114, 397–407 (2004).

34. Horii M, Touma O, Bui T, Parast MM. Modeling human trophoblast, the placental epithelium at the maternal fetal interface. Reproduction 160, R1–R11 (2020).

35. Turco MY, Moffett A. Development of the human placenta. Development 146, dev163428 (2019).

36. Lee CQ, et al. What is trophoblast? A combination of criteria define human first-trimester trophoblast. Stem cell reports 6, 257–272 (2016).

37. Malassiné A, Frendo JL, Evain-Brion D. A comparison of placental development and endocrine functions between the human and mouse model. Hum Reprod Update 9, 531–539 (2003).

38. Sheridan MA, et al. Establishment and differentiation of long-term trophoblast organoid cultures from the human placenta. Nature Protocols 15, 3441–3463 (2020).

39. Lew RA, et al. Angiotensin-converting enzyme 2 catalytic activity in human plasma is masked by an endogenous inhibitor. Experimental Physiology 93, 685–693 (2008).

40. Lambert DW, et al. Tumor necrosis factor-α convertase (ADAM17) mediates regulated ectodomain shedding of the severe-acute respiratory syndrome-coronavirus (SARS-CoV) receptor, angiotensin-converting enzyme-2 (ACE2). Journal of Biological Chemistry 280, 30113–30119 (2005).

41. Lumbers E, Delforce SJ, Arthurs AL, Pringle KG. Causes and Consequences of the Dysregulated Maternal Renin-Angiotensin System in Preeclampsia. Frontiers in Endocrinology 10, (2019).

42. Tukiainen T, et al. Landscape of X chromosome inactivation across human tissues. Nature 550, 244–248 (2017).

43. Sykes S, Pringle K, Zhou A, Dekker G, Roberts C, Lumbers E. The balance between human maternal plasma angiotensin II and I angiotensin 1-7 levels in early gestation pregnancy is influenced by fetal sex. Journal of the Renin-Angiotensin-Aldosterone System 15, 523–531 (2014).

44. Melamed N, Yogev Y, Glezerman M. Fetal gender and pregnancy outcome. The Journal of Maternal-Fetal & Neonatal Medicine 23, 338–344 (2010).

45. Di Renzo GC, Rosati A, Sarti RD, Cruciani L, Cutuli AM. Does fetal sex affect pregnancy outcome? Gender medicine 4, 19–30 (2007).

